# Evolution of endogenized densoviral elements across aphid species

**DOI:** 10.64898/2026.07.29.741575

**Authors:** Paula Rozo-Lopez, Joseph Torres, Vanesa Torres, Meaghan J. Adler, Keertana Tallapragada, Oliver Cope, Benjamin J. Parker

## Abstract

Endogenous viral elements (EVEs) are widespread across animal genomes, yet the processes governing EVE evolution and diversification remain poorly understood. Here, we characterize the evolution of densoviral EVEs and exogenous densoviruses across the aphid tribe Macrosiphini, an agriculturally important group in which exogenous densoviruses and their endogenous derivatives have been linked to the plastic production of wings. Using new genome assemblies, transcriptomics, and phylogenetic analysis, we find that EVE content varies extensively across species. Moreover, we discovered a novel densovirus that is vertically transmitted, but phylogenetic incongruence between other densoviruses and their hosts suggests that horizontal transmission may also occur. Finally, we show that EVE-mediated regulation of wing plasticity extends across species that use different environmental signals to induce winged offspring. Our study shows that in this system, the evolution of EVEs is highly variable and lineage-specific, generating genomic patterns that cannot be predicted from host evolutionary relationships.

## Introduction

Viral sequences integrated into host genomes, known as endogenous viral elements (EVEs), have been identified across the tree of life (Katzourakis and Gifford 2010). Viruses interact closely with host cellular machinery, and the sequences they leave behind upon endogenization can be expressed and co-opted by hosts (Blair, et al. 2020; Frank and Feschotte 2017; Ter Horst, et al. 2019). EVEs have been implicated in innate immunity, development, and gene regulation (Holmes 2011), suggesting that co-option is a recurring feature of genome evolution.

In recent years, it has become clear that many viruses that form heritable symbioses with their hosts are vertically transmitted from parents to offspring (Parker and Rozo-Lopez 2026). These symbioses are maintained, in part, because they shape host phenotypes in ways that ensure their own transmission, conferring benefits such as protection against natural enemies, while also manipulating host reproduction, behavior, and interactions with other species (Coffman, et al. 2022; Kageyama, et al. 2023; Nagamine, et al. 2023; Ryabov, et al. 2009; Varaldi, et al. 2023; Wang, et al. 2017; Xiao, et al. 2021). Vertical transmission also means that viruses are in contact with host germlines across generations, providing repeated opportunities for endogenization.

Interactions between aphids and densoviruses, single-stranded DNA densoviruses in the family *Parvoviridae*, provide a unique system for studying both heritable symbiosis and EVEs. Both exogenous densoviruses (Guo, et al. 2022; Li, et al. 2022; Li, et al. 2025; Pinheiro, et al. 2019; Ryabov, et al. 2009; Tang, et al. 2016; van Munster, et al. 2003a; van Munster, et al. 2003b) and their endogenous derivatives (Clavijo, et al. 2016) are present across the aphid tribe Macrosiphini, an agriculturally relevant group of aphids. Importantly, this system has a well characterized phenotype: some densoviruses manipulate the environmentally-induced production of winged or wingless offspring (Dong, et al. 2024; Pinheiro, et al. 2019; Ryabov, et al. 2009), which is a key life history trait (Yang and Pospisilik 2019). This manipulation presumably enhances viral dispersal and transmission (Rozo-Lopez and Parker 2023). Strikingly, in pea aphids, endogenized densoviral nonstructural proteins have been co-opted to regulate wing plasticity (Parker and Brisson 2019). Densoviral EVEs have also been shown to be transcriptionally active in other species (Clavijo, et al. 2016); however, whether this co-option is conserved in other aphid species, and how densoviral EVEs have evolved across the clade, remains unknown.

In this study, we characterized the distribution of densoviral EVEs and exogenous densoviruses across the aphid tribe Macrosiphini, integrating new aphid genome assemblies, a newly discovered densovirus, transcriptomic data, and phylogenetic analyses. We show that densoviral EVEs are widespread and vary extensively among closely related species without a clear phylogenetic pattern, suggesting high genomic turnover and/or repeated endogenization. Furthermore, we find that viral infections and EVE presence are not consistently associated across host lineages, and that exogenous densoviruses show evidence of horizontal transmission across clades. Finally, we show that co-opted viral genes are expressed in association with the wing-induction cue in species that rely on different environmental triggers, consistent with conserved co-option. Together, our results demonstrate that endogenization of viral symbionts can drive rapid, lineage-specific evolution in aphids.

## Results

### Densoviral sequences are widespread among aphid transcriptomes

To characterize expressed densoviral sequences across the Macrosiphini, we field-collected aphids from 11 species and maintained them as clonal laboratory lines for at least three generations before sampling RNA. We screened these samples with CZ ID (Simmonds, et al. 2024) and detected densoviral reads in 8 of the 11 species (Fig. 1a). We then used TRAVIS, a consistency-based virus detection pipeline, to conduct a more extensive screen of our dataset for Parvoviridae proteins with greater than 60% similarity to densoviral ORFs, identifying 642 densoviral-like contigs (500–1098 nt). We then screened existing RNAseq datasets from the Sequence Read Archive (Table S1), and identified densoviral reads (at a cutoff of 0.15 NT rPM) in 8 of 25 Macrosiphini species evaluated (Fig. 1b). Together, these analyses showed that densoviral sequences, whether from exogenous viruses, transcribed EVEs, or both, are a common feature of Macrosiphini transcriptomes.

**Figure 1.**
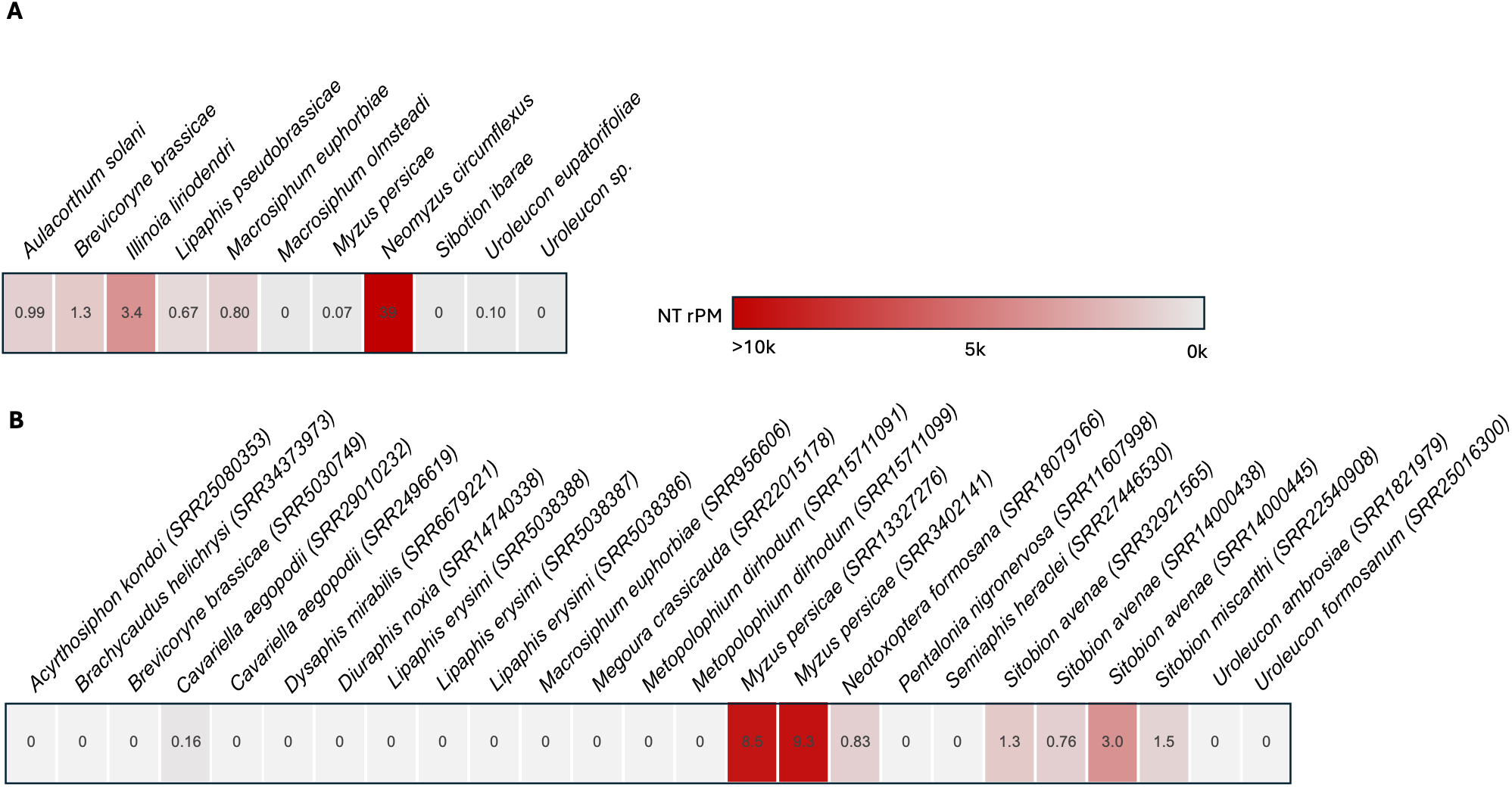
Relative abundance of densoviral hits from RNAseq of (**a**) field-collected aphids and (**b**) SRA datasets from different aphid species. The NT rPM gradient bar indicates the proportion of reads aligning to densoviruses in the NCBI nucleotide database, with specific values shown in parentheses. RNAseq accession numbers are listed in Table S1.

### Macrosiphini genome sequencing and phylogenetics

To distinguish exogenous viral infections from transcribed EVEs, and to provide genomic resources for downstream analyses, we assembled the genomes of two key Macrosiphini species. Our assembly of *Uroleucon eupatorifoliae* is 452 Mb with 29.5% GC content, an N50 of 1.58 Mb, 99.1% BUSCO completeness, and 10.3% duplication (ASM4999865v1; Fig. S1). Our assembly of a genome for *Neomyzus circumflexus* is 391 Mb with 30.3% GC content, has an N50 of 2.15 Mb with 98.4% BUSCO completeness, and 1.3% duplication (ASM5896979v1; Fig. S1). These genome sizes and GC contents fall within the range of other aphid genomes sequenced to date (Shigenobu and Yorimoto 2022).

Determining the phylogenetic relationships among Macrosiphini has been challenging due to the absence of shared morphological traits and varying results from different genetic markers, which produce conflicting associations among some groups (Choi, et al. 2018; von Dohlen, et al. 2006).

Therefore, to reconstruct the evolutionary relationships among Macrosiphini species, we obtained ultra-conserved elements (UCEs) from each species with a sequenced genome and we constructed a maximum likelihood phylogeny (Fig. 2a, Fig. S2). Our UCE-based phylogeny differs from previous studies (Choi, et al. 2018) in several ways: it places the model aphid *A. pisum* as an outgroup to a clade containing *Sitobion* and *Macrosiphum*; it places *Neomyzus* as a sister clade to the genus *Uroleucon*; and it places *Brevicoryne brassicae* as the outgroup of a clade containing *Diuraphis* and *Semiaphis*, rather than *Diuraphis* as the outgroup. These new genomes and the resulting phylogenetic framework provide a necessary foundation for comparative studies within the Macrosiphini.

**Figure 2.**
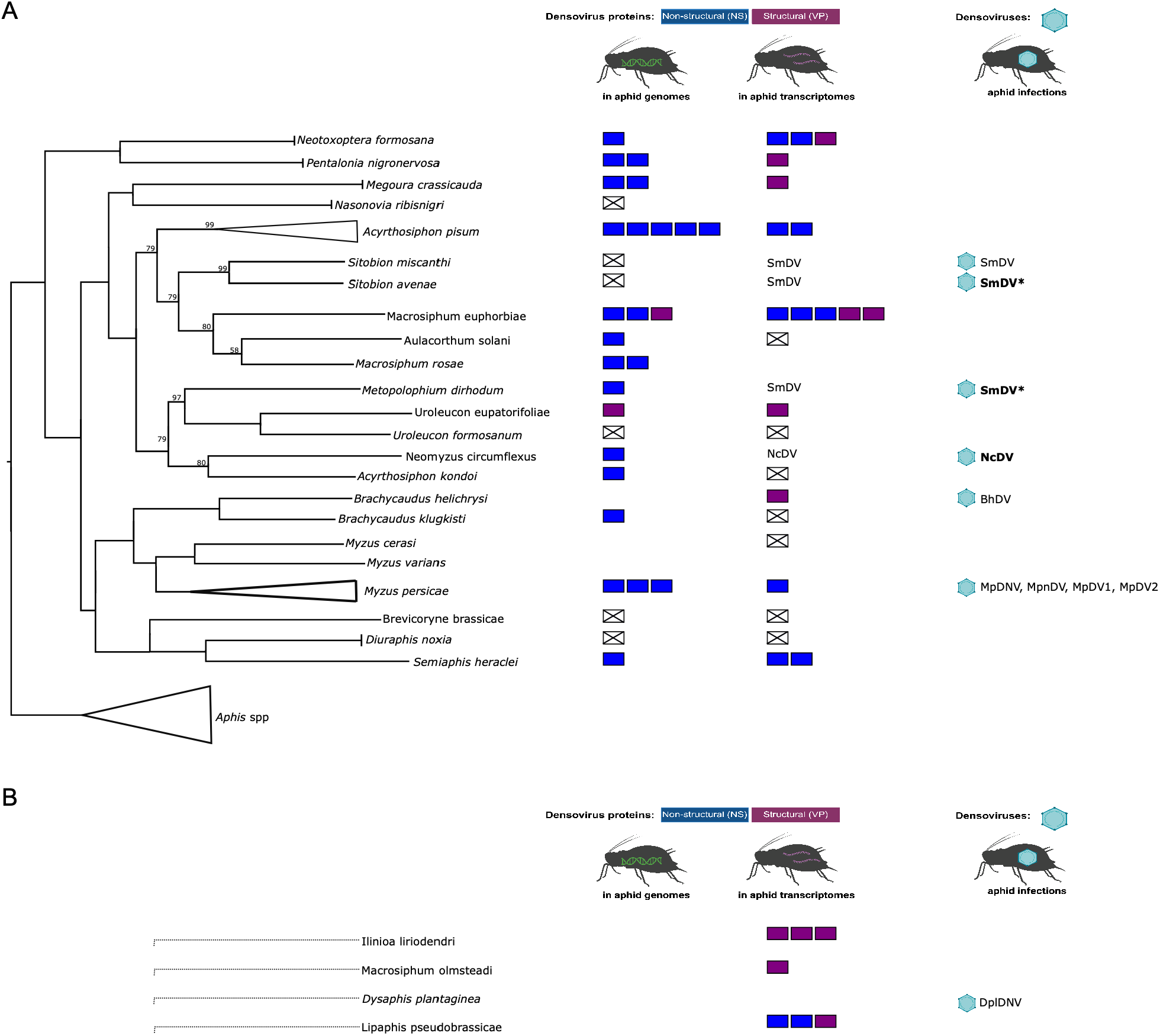
Densoviral findings across Macrosiphini aphids. (**a**) Phylogenetic representation (left) of Macrosiphini based on a maximum-likelihood analysis of UCEs from representative aphid genomes (NCBI and BIPAA) (Figure S2). Bootstrap support values below 100 are shown on the corresponding branches and represent percentages from 500 replicates. The tree was inferred unrooted with RAxML-NG and rooted for display using Aphis as the outgroup. EVE densoviral findings from NCBI datasets (middle) are shown as colored boxes (blue non-structural protein-like and purple structural protein-like sequences) to indicate the number of densoviral EVE copies in aphid genomes and those that are actively transcribed. White boxes with an “X” indicate the absence of densoviral EVE sequences. Aphid species without genomes or transcriptome datasets available in NCBI were left as empty spaces (Table S1). Exogenous densoviruses (right) are depicted as a virus capsid followed by the virus species acronym. (**b**) Macrosiphini species for which phylogenetic placement could not be determined due to the lack of a reference genome.

### Densoviral EVEs are variable and evolutionarily dynamic

We screened for densoviral hits (either EVEs or potential densoviral infections) across the aphid phylogeny using available NCBI genomic data (Table S1). We then used the transcriptomic data described above to determine whether the genomic origin of actively transcribed EVEs corresponded to viral insertions encoding structural (VP) or non-structural (NS) densoviral proteins. Our analysis (Fig. 2) revealed repeated, widespread viral endogenization of sequences with high homology to densoviral NS proteins, which predominate across the phylogeny and are actively transcribed in many aphid species. We also found that EVE copy number varies substantially across species, with some lineages harboring up to five copies (in the *A. pisum* genome) but most harboring only a few (one to three). This variation is not explained by phylogenetic position alone, which is inconsistent with a single ancient endogenization; the pattern could instead reflect ongoing, independent integration events, lineage-specific duplication and loss of a smaller number of ancestral insertions, or some combination of these processes. Together, these patterns reveal a genomic landscape that is neither static nor phylogenetically predictable.

### Exogenous densoviruses are distributed across aphid clades and show evidence of host switching

In addition to previously reported aphid densoviruses: SmDV (Li, et al. 2022), BhDV (Li, et al. 2025), MpDV (van Munster, et al. 2003a; van Munster, et al. 2003b), MpnDV (Tang, et al. 2016), MpDV1 (Sukhikh, et al. 2024), MpDV2 (Guo, et al. 2024), and DplDV (Ryabov, et al. 2009), we further characterized the diversity and evolutionary history of exogenous densoviruses across Macrosiphini. We first determined whether densoviral hits in our transcriptomic data represented actively transcribed EVEs or circulating viruses (Fig. 2). In *Neomyzus circumflexus*, initial screening revealed assembled contigs (3900–6900 nt) with less than 70% similarity to any known densovirus. Crucially, no full-length matches to these contigs were found in the *N. circumflexus* genome assembly, suggesting they did not derive from integrated EVEs. Using these contigs to scaffold nanopore DNA reads with at least 1kb overlap, 99% identity, and with read lengths close to the expected densoviral genome size (5.5–6 kb), we assembled a consensus viral genome of 5947 nt from combined RNA and DNA reads (Fig. 3a). Using the NCBI web version of ORFfinder, we identified open reading frames corresponding to the major non-structural (NS1, 1887 nt; NS2, 987 nt) and structural (VP1, 2166 nt; VP2, 654 nt) proteins (Fig. 3b). Based on sequence-identity criteria for parvovirus classification (Penzes, et al. 2020), we described this as a new species, *Neomyzus circumflexus* densovirus (NcDV), and deposited the reference genome in NCBI (PZ554439). To assess heritability, we PCR-tested individual adults from the field-collected parental generation (P) and from at least three subsequent laboratory generations (F1 through F3), a key criterion for determining vertical transmission (Parker and Rozo-Lopez 2026). NcDV was maintained with 100% fidelity across 26 generations of laboratory colonization (one year), consistent with heritable vertical transmission.

**Figure 3.**
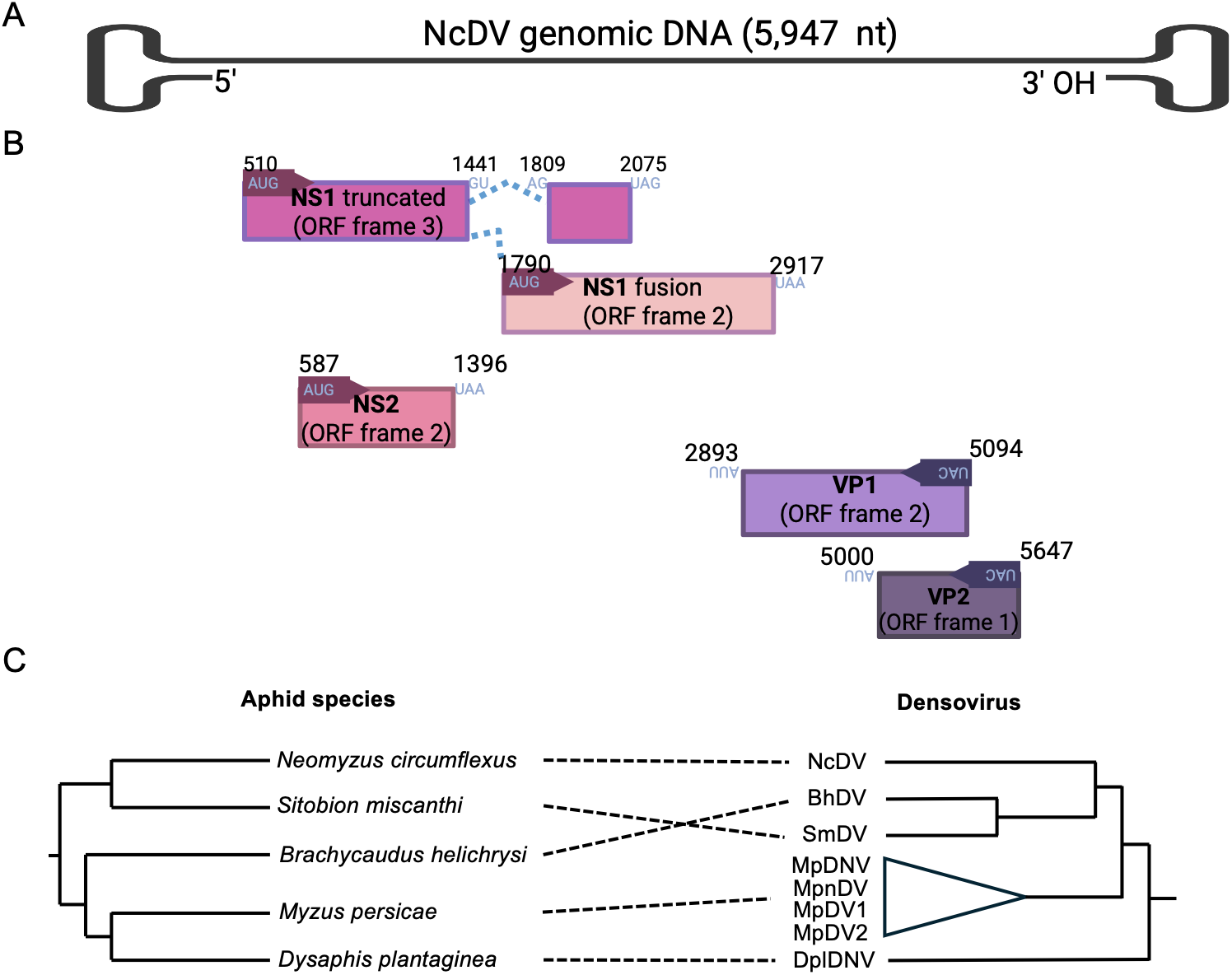
(**a**) Genome and **(b)** transcript organization of *Neomyzus circumflexus densovirus* (NcDV) revealed by *de novo* assembly of Illumina (RNA-seq) and Nanopore (DNA) reads from *N. circumflexus* aphids. The viral genomic ssDNA (5,947 nt) is depicted in black, with viral transcripts shown as pink and purple boxes. Major transcription start sites (arrows) of the rightward non-structural (NS) and leftward capsid protein (VP) genes are shown with pink and purple bent arrows, respectively. Spliced mRNAs transcribed from the NS units are shown with blue and red bent arrows. The encoded proteins are labeled in bold (inside the boxes), and the ORFs’ start and stop codons (blue font) and sites (black numbers) are labeled above the boxes. The ORF frame shift is produced by leaky scanning. (**c**) Representation of phylogenetic trees of aphid species (left) and aphid densoviruses (right) with dotted lines indicating which hosts harbor infections of a particular densovirus.

We also performed a combined analysis of NCBI genomic and transcriptomic reads (Fig. 2a) that had BLAST-matched regions with at least 1kb overlap and 99% identity to existing densoviral genomes, resulting in the assembly of five densoviral genomes (Table S2). Overall, we observed that the distribution of EVEs does not mirror that of all exogenous densoviruses in our aphid phylogeny (Fig. 2). For example, species harboring EVEs are not consistently those with active viral infections, and vice versa, demonstrating that EVEs and exogenous viruses can persist and be lost independently across host lineages.

To examine the broader relationship between densovirus and aphid phylogenies, we constructed a phylogeny of all known aphid densoviruses using NS1 amino acid sequences (Cotmore, et al. 2019) and compared it with the Macrosiphini host phylogeny (Fig. 3c). We found that densoviruses are distributed across multiple aphid clades with no single lineage monopolizing infection. SmDV, for example, was detected in three distantly related species: *Sitobion avenae, S. miscanthi*, and *Metopolophium dirhodum* (Fig. 3c). The densovirus phylogeny shows clear incongruence with the host phylogeny, as closely related viruses infect distantly related aphid species. This pattern is inconsistent with strict vertical transmission and instead points to horizontal transmission and host switching across aphid lineages over evolutionary time.

### Differential expression of a densoviral gene in a second aphid species

In the pea aphid (*Acyrthosiphon pisum*), densoviral EVEs have been shown to regulate wing plasticity, with two APNS genes upregulated in response to crowding and required for normal wing induction (Parker and Brisson 2019). To test whether this functional role extends beyond the pea aphid, we examined wing plasticity and densoviral EVE expression in three Macrosiphini species: *Macrosiphum euphorbiae, Aulacorthum solani*, and *Illinoia liriodendri*. Crowding did not significantly increase the proportion of winged offspring in any of the three species relative to solitary conditions (Fig. 4a), indicating that the crowding response observed in pea aphids is not universal across Macrosiphini. However, we found that *M. euphorbiae* adults maintained on low-quality host plants produced a higher proportion of winged offspring than those on high-quality plants (Fig. 4b), demonstrating that wing plasticity in this species responds to a different environmental cue (host plant quality rather than crowding). The *M. euphorbiae* genome contains an Apns-2 homolog with close sequence similarity to the pea aphid densoviral insertion (Rozo-Lopez, et al. 2023), which was incorrectly described as a novel densovirus (Teixeira, et al. 2018).

**Figure 4.**
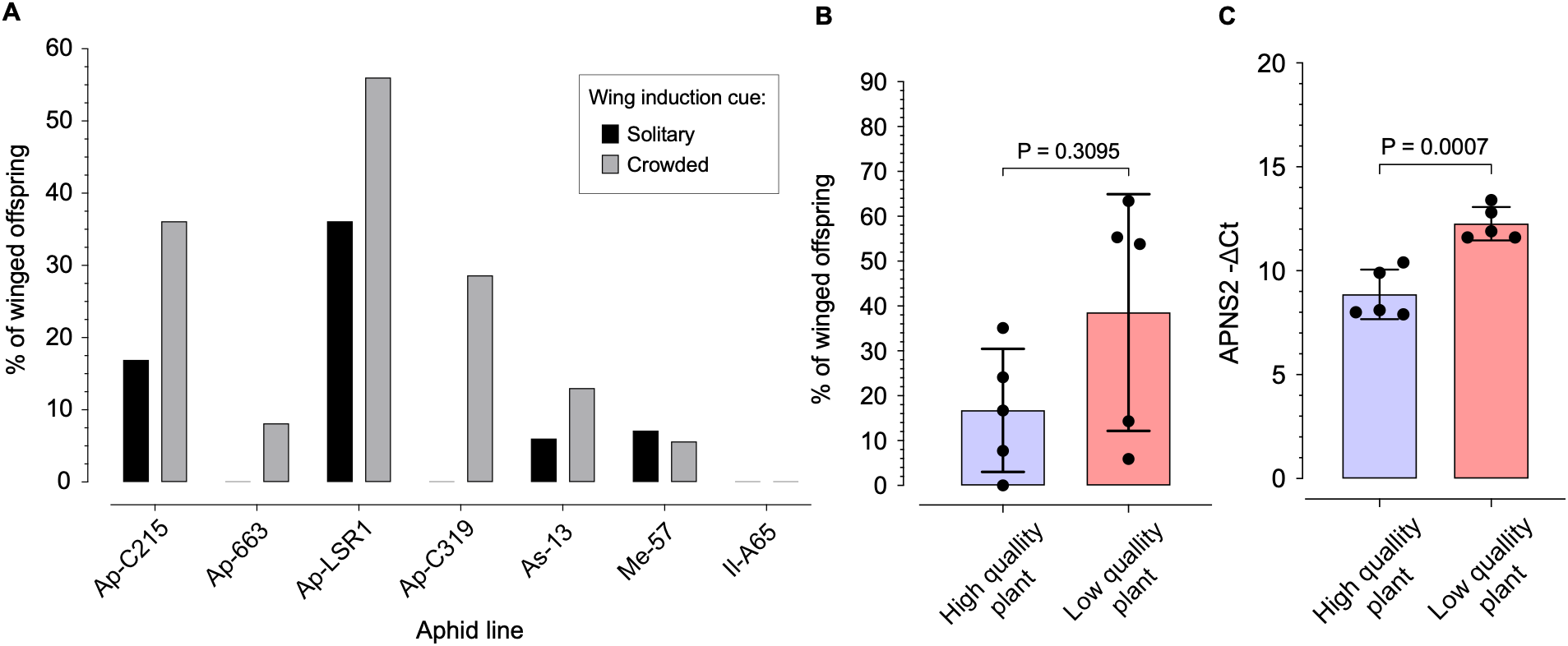
Wing induction and densoviral gene expression. **(a)** Percentage of winged offspring (y axis) produced when aphid lines are exposed to solitary or crowded conditions (*A. pisum*: Ap; *A. solani*: As; *M. euphorbiae*: Me; *I. liriodendri*: Il). **(b)** Percentage of winged offspring produced by *M. euphorbiae* parental aphids reared on high and low quality tomato plants. **(c)** Change in gene expression (qPCR) of the *M. euphorbiae* APNS homolog in parental aphids producing wingless and winged offspring in response to high and low plant quality conditions, respectively. **(b and c)** Each point represents a pool of four adult aphids. A Mann-Whitney test was used for statistical evaluation.

Using qRT-PCR, we found that this gene is significantly upregulated in adults producing winged offspring in response to low plant quality, compared with adults maintained on high-quality plants (p = 0.0007, Fig. 4c). This mirrors the upregulation of APNS genes during wing induction in pea aphids despite a different environmental trigger. Together, these results suggest that the functional co-option of densoviral NS genes for wing plasticity is not unique to the pea aphid but extends to at least one additional Macrosiphini species. Crucially, this co-option persists even when the environmental context triggering wing production differs between species, suggesting that the downstream regulatory role of these EVEs is conserved even as the upstream induction pathway has diverged.

## Discussion

Aphid densoviruses provide an opportunity to study endogenous viral elements and their homologous viruses simultaneously. To enable this work, we sequenced two new Macrosiphini genomes and used them to revise the phylogenetic framework for the group, providing the resolution needed to distinguish exogenous viral infections from transcribed EVEs and to examine their distribution across the phylogeny. We show that densoviral EVEs are widespread across Macrosiphini, but their distribution and copy number lack a strong phylogenetic pattern, suggesting that integration, duplication, and/or loss are frequent lineage-specific events rather than the result of an ancient endogenization. We also discovered a new densovirus with evidence of strong vertical transmission among *Neomyzus circumflexus* aphids. Our analysis shows that exogenous densoviruses exhibit incongruence with the host phylogeny, consistent with horizontal transmission rather than strict vertical co-divergence.

How co-opted APNS genes function mechanistically remains an open question. In pea aphids, crowding-induced wing production is associated with upregulation of APNS genes (Parker and Brisson 2019). Our results extend this functional relevance to *M. euphorbiae*, where we found that densoviral gene upregulation responds to host plant quality. This result suggests that APNS gene function in wing induction is conserved even as the upstream environmental trigger differs between species. Another key unresolved question is whether the function of EVEs reflects the function of the exogenous virus during active infection. A recent paper found that MpDNV-positive aphids produce elevated levels of the alarm pheromone (E)-farnesene (Dong, et al. 2024), which is a known inducer of wing production (Wang, et al. 2025). Future work is needed to determine whether APNS proteins act through the same mechanism, via functional experiments on alarm pheromone production and sensitivity.

Beyond their co-option for developmental functions, EVEs derived from non-retroviruses have been shown in other insects to serve as templates for piRNA-mediated antiviral immunity, with EVE-derived small RNAs silencing homologous exogenous viruses (Ter Horst, et al. 2019). Whether aphid densoviral EVEs serve an analogous immune function remains unknown. In *M. persicae*, MpDNV interacts with the aphid RNAi pathway; the silencing suppressor activity of the plant viral protein PLRV P0 elevates MpDNV titer, suggesting that siRNA-based immunity normally suppresses densoviral replication in aphids (Preising, et al. 2026); however, it was unclear from this study whether EVEs play a role in viral silencing. We found that several aphid species harboring exogenous densoviruses also have densoviral EVEs, but SmDV infects hosts that appear to lack corresponding densoviral EVEs.

This absence is notable given that SmDV has been described as measurably pathogenic (Li, et al. 2022) and, based on our phylogenetic analysis, appears to infect multiple distantly related host species. This pattern is consistent with the possibility that EVEs contribute to the stability of densoviral heritable symbiosis (Clavijo, et al. 2016), but further data are needed.

Our results suggest that heritable viral symbioses challenge the traditional view of non-retroviral EVEs as rare, ancient integration events. Instead, the densoviral EVE landscape in Macrosiphini appears to be lineage-specific and rapidly evolving, driven by ongoing integration, duplication, and loss rather than by a small number of ancient events. Many aphid genomes contain numerous EVEs related to other viruses (Liu, et al. 2020), and in general, insect genomes have a high density of EVEs (Ter Horst, et al. 2019). As the diversity of heritable viral symbioses becomes more apparent across invertebrates, heritability should be considered a key variable for predicting EVE evolution and interpreting the genomic signatures they leave behind.

## Materials and methods

### Aphid collection and colonization

We collected adult winged and wingless asexual female aphids infesting different host plant species across urban and agricultural landscapes in East Tennessee, USA, during April–June and October–December 2022–2024. We placed single field-collected aphids in Petri dishes containing a leaf disc of the appropriate host plant and allowed them to reach parthenogenetic reproduction in the laboratory. Then, we collected surviving parental adults that had produced at least five offspring and stored each parental aphid individually in Eppendorf tubes at −80ºC for further RNA extraction and sequencing. We subsequently used the offspring to establish laboratory colonies and maintained them on the respective host plants (Table S1) at 20ºC 16L:8D. We identified each aphid species as previously reported (Foottit, et al. 2008; Rozo-Lopez, et al. 2023) by DNA extraction, PCR amplification, and Sanger sequencing of the COI barcode region.

### Metatranscriptome sequencing

We macerated frozen aphids with a pestle in 300 µL of TRIzol (Invitrogen; Thermo Fisher Scientific, Inc., Waltham, MA, USA) with 100 μL BCP (1-bromo-3-chloropropane; Life Technologies, Thermo Fisher Scientific, Inc., Waltham, MA, USA) and isopropanol precipitation. We resuspended each RNA pellet in 40 μL nuclease-free water, removed genomic DNA using Zymo DNase I Reaction Mix according to the manufacturer’s instructions (Zymo Research, Irvine, CA), and cleaned and concentrated the RNA with the Zymo RNA Clean & Concentrator kit under recommended conditions. We pooled equal amounts of RNA from 1–6 aphids of the same species (Table S3) to perform metatranscriptome sequencing at Novogene (Novogene Corporation Inc., Sacramento, CA, USA). Library preparation was conducted using the Illumina TruSeq Stranded Total RNA with Ribo-Zero Plus and NEBNext rRNA Depletion Kit. Libraries were sequenced to approximately 9 billion base pairs (bp) per sample with 150 bp paired-end reads on an Illumina NovaSeq platform. Raw reads were deposited in the NCBI Sequence Read Archive (Table S1).

### *Densovirus* detection in RNAseq data

We used the CZ ID platform (https://czid.org) with Illumina mNGS Pipeline v8.2 (Kalantar, et al. 2020; Zendesk 2023) as previously described (Rozo-Lopez, et al. 2023). In addition, we screened for Densovirus-like open reading frames (ORFs) using TRAVIS (v.20221029, https://github.com/kaefers/travis) and the 2022 Densovirinae reference database (Rozo-Lopez, et al. 2023) on *de novo* transcriptome assemblies generated with Trinity v.2.15.1 using default settings (Grabherr, et al. 2011). Densovirus-like ORFs (100–2000 amino acids) were extracted from the assembled transcriptomes and screened using HMMER v3.3.1 (Wheeler and Eddy 2013), MMSeqs2 (Steinegger and Söding 2017), BLASTP v2.12.0 (Altschul, et al. 1997), and Diamond v2.0.15 (Buchfink, et al. 2021) with an e-value cutoff of 1×10^-6^. *M. euphorbiae* (Rozo-Lopez, et al. 2023) and *A. pisum* (Parker and Brisson 2019) screening was based on previously published analyses.

### *Uroleucon eupatorifoliae* DNA extraction and sequencing

We pooled five genetically identical adult unwinged aphids from the cultivated lab line A120 (Table S3) and isolated genomic DNA (gDNA) using Bender buffer and ethanol precipitation (Bender, et al. 1983). We resuspended the DNA pellet in 50 μL of nuclease-free water, removed RNA using the Zymo RNase A reaction mix according to the manufacturer’s instructions, and cleaned and concentrated the DNA with the Zymo DNA Clean & Concentrator kit under recommended conditions. For library preparation, we used 1 μg of gDNA, the Nanopore Ligation Sequencing Kit SQK-LSK110 (Oxford Nanopore Technologies, UK), and the NEB Companion Module (New England Biolabs, Ipswich, MA, USA), following the manufacturer’s recommendations. We used a Nanopore R.9.4.1 (FLO-MIN106D) flow cell with a MinION Mk1B (MIN 101-B) sequencing device (Oxford Nanopore Technologies, Oxford, UK). We sequenced the libraries for 24 hours on the flow cell with base calling turned off.

### *Neomyzus circumflexus* DNA extraction and sequencing

High molecular weight (HMW) gDNA from *Neomyzus circumflexus* (line B045; Table S3) was extracted using Qiagen Genomic-tip 20/G columns (Qiagen, Hilden, Germany) with an adapted version of the user-developed protocol, “Isolation of genomic DNA from mosquitoes or other insects using the QIAGEN Genomic-tip-(EN)”. Two aphids per sample were pestle-homogenized on dry ice in a 1.5 mL DNA LoBind microcentrifuge tube, then suspended in 1 mL of a custom lysis buffer (per 100 mL: 99 mL Qiagen AE Elution Buffer, 0.585 g NaCl, 4.78 g Guanidine-HCl, 1 mL Triton X-100 (v/v)). Lysates were treated with RNase A (0.1 mg/mL) at 37°C for 30 min, followed by Proteinase K (0.8 mg/mL) at 50°C for 3 h with gentle agitation. After centrifugation (15,000 x g for 20 min, 4°C), the supernatant was incubated at 4°C for 12 h to enhance protein precipitation, then centrifuged again and loaded onto an equilibrated Qiagen Genomic-tip 20/G. Genomic-tip wash steps followed the manufacturer’s protocol; DNA was eluted twice with 800 μL of 50°C Buffer QF. Eluates were precipitated with 600 μL isopropanol and glycogen (0.02 mg/mL), mixed overnight at room temperature on a HulaMixer, then pelleted (15,000 x g for 20 min, 4°C), washed twice with 80% ethanol, and resuspended in TE buffer (10 mM Tris, pH 8.5, 0.1 mM EDTA) at 4°C for 2 days with mixing. DNA quality and quantity were assessed using a NanoDrop, a Qubit, and a TapeStation 4200 (Agilent Technologies Inc., Santa Clara, CA). Samples were pooled totaling 8 input aphids and sequenced by Novogene Inc. on the Nanopore PromethION DNA library platform.

### Aphid genome assembly

We first evaluated raw data with Jellyfish (Marçais and Kingsford 2011) and Genomescope2 (Ranallo-Benavidez, et al. 2020) to estimate the genome size to pass onto the assembler with a k-mer histogram of length 21. We assembled *Uroleucon* Nanopore reads with NextDenovo (Hu, et al. 2024) with the recommended default parameters and an estimated genome size of 450 Mb. We assembled Neomyzus Nanopore reads with NextDenovo using the recommended default parameters and an estimated genome size of 384 Mb. After assembly, we removed all contigs that matched a phylum other than Arthropoda, including those from Proteobacteria, using the BlobToolKit visualizer (Challis, et al. 2020) after searching the contigs against the NCBI nucleotide NR database with the BLAST+ (Camacho, et al. 2009) command-line interface. We evaluated the N50 and measured completeness using BUSCO v6.0.0 (Manni, et al. 2021) with the default parameters and the appropriate Hemiptera lineage. Gene predictions were then carried out on the assembly using Helixer (Stiehler, et al. 2020) with the provided invertebrate model. The resulting gene predictions were then used for automated functional annotation with the EnTAP pipeline (Hart, et al. 2020) using the default eggNOG-mapper with homology searches against the UniProt TrEMBL database, NCBI NR database, Ref-seq invertebrate database, and the Swiss-Prot database. Homology information was then incorporated into the GFF file using AGAT (Dainat, et al. 2025). The NCBI genome accession numbers are available in Table S1 and the bioinformatic code is available at https://github.com/Artifice120/Uroleucon-eupatorifoliae-assembly-to-annotation and https://github.com/Artifice120/Neomyzus-assembly

### Macrosiphini phylogeny

In addition to the aphid genomes produced in-house, we downloaded all publicly available reference genomes (NCBI and BIPAA AphidBase), as of January 2026, for aphid species belonging to the tribe Macrosiphini (Table S1). We captured ultra-conserved elements (UCEs) from each genome using the phyluce v1.7.3 workflow, as previously described for Hymenoptera UCEs (Branstetter, et al. 2017). Then, we used OrthoFinder v2.5.5 (Emms and Kelly 2019) to cluster the UCEs into 189 orthogroups representing conserved sequences descended from a common ancestor, with multiple sequence alignments performed via all-versus-all comparisons of sequences, requiring that at least 80.5% of species have single-copy genes in any orthogroup. In the OrthoFinder pipeline, MAFFT with default settings was used to generate the initial multiple sequence alignments (MSAs), and a species tree was generated using the STAG algorithm (Emms and Kelly 2018) as an intermediate step. The concatenated species-tree alignment produced by OrthoFinder was then tested to find the best-fit model using ModelTest-NG (Darriba, et al. 2019) with the transversional model and a gamma distribution, with four categories being identified as the best fit. Finally, we used this concatenated alignment to construct a phylogenetic tree (shown in Figure 2) with RAxML-NG v2.0.1 (Kozlov, et al. 2019) using the TVM+I+G4 substitution model. Phylogenetic inference was performed with 500 bootstrap replicates. The tree was inferred unrooted; UCEs from aphids of the genus Aphis were assigned as the outgroup for display (Table S4). Bioinformatic code is available at https://github.com/Artifice120/UCE_Aphidae_Phylogeny-

### Characterizing endogenous viral elements in aphid genomes

Raw DNA nanopore reads from *U. eupatorifoliae* and *N. circumflexus* were used as input to the CZ ID platform pipeline V0.7 (https://czid.org) with a z-score ≥1 and NT L ≥50, as described above (Bohl, et al. 2022; Kalantar, et al. 2020). We conducted genomic screening for densoviral EVEs across the 20 genomes available in NCBI (Table S1) by performing discontinuous megablast searches of the major aphid densoviral proteins and the densoviral EVEs (Table S5), using an expect threshold of 0.05, linear gap costs, a match score of 1, and a mismatch score of −2 with no filtering. We then retained non-redundant hits with E-values < 1×10^-3^ and a BLAST-matched region of at least 85% identity to the query. All matches to specific densoviral proteins were confirmed with ORFfinder.

### *Neomyzus circumflexus densovirus* (NcDV)

We downloaded CZ ID reads (RNAseq) matching *Ambidensovirus* and non-genus-specific reads in the *Parvoviridae* family, and assembled them *de novo* in Geneious at medium sensitivity (Geneious Prime v.2023.2.1). We selected viral contigs longer than 2,000 nt. We aligned the CZ ID contigs with the viral hits (>1,000 nt) provided by TRAVIS using the Geneious Prime Clustal Omega v.1.2.3 plugin and the mBed algorithm. We selected alignments with pairwise identity greater than 98% and BLAST homology greater than 70% to other densoviruses (BhDV & MpDV2) as a partial consensus sequence. The partial genome sequence (5,442 nt) generated from this was used as a query to search for individual Nanopore reads ranging from 2–6 kb. We selected reads with a BLAST match region of at least 1 kb overlap and 99% identity to our query, then used these to build a new assembly with Canu v2.2. We used the NCBI web version of ORFfinder with the standard genetic code to identify viral proteins. The NcDV consensus genome is available on NCBI with accession PZ554439.

### NcDV PCR

We designed primers to detect NcDV in aphids (NcDV_4001F 5’– GCTTCCGAAATTGTTCCTGC–3’ and NcDV_4507R 5’–TGAGTTACATACGCACTGGC–3’). We confirmed primer specificity by retesting several aphid species with known densoviral EVEs (based on

RNAseq analysis). For initial NcDV screening, we used 800 ng of total RNA (extracted as described for RNAseq) for cDNA synthesis using the iScript cDNA synthesis kit (Bio-Rad Laboratories, Inc., Hercules, CA, USA), which employs random hexamer primers. Either 1 μL of undiluted cDNA or 80 ng of gDNA, extracted with Bender buffer (as above), served as templates in 25 μL PCR reactions using Quick-Load Taq 2X Master Mix (New England Biolabs, Ipswich, MA, USA), 0.2 μM of each primer, and the following thermal cycling conditions: 94ºC for 5 min, then 35 cycles of 94ºC for 30 s, 60ºC for 30 s, 72ºC for 30 s, and a final extension at 72ºC for 3 min. PCR products were visualized on 1% agarose gels stained with SYBR Safe (Invitrogen, Inc.).

### Densovirus phylogeny

We aligned the non-structural protein 1 (NS1) sequences (Cotmore, et al. 2019) of newly and previously reported aphid densoviruses (Table S6) in MAFFT v7.490 using the FFT-NS-i_x1000 algorithm, matrix BLOSUM62, and 1.54 gap penalty (Katoh, et al. 2002). Then, we used the amino acid alignment to generate a maximum–likelihood tree in RAxML v8.2.12 with the PROTGAMMAJTTF model and a majority-rule bootstrap with 500 replicates (Stamatakis 2014). We used the NS1 sequence of *Galleria mellonella densovirus* (GmDNV, NC_004286.1) as the outgroup.

### Wing induction assays

We used colonized *Macrosiphum euphorbiae* (line Me57) maintained on tomato plants (Husky Cherry Red), *Illinoia liriodendri* (line A65) maintained on shepherd’s purse (*Capsella bursa-pastoris*), and *Aulacorthum solani* (line As13) and *Acyrthosiphon pisum* (lines C215, 663, LSR1, C319) maintained on fava plants (Vroma). All aphid lines were reared at low densities at 20°C under 16L:8D. Initially, we exposed stage-four nymphs to crowding (12 aphids in a Petri dish) and solitary conditions (1 aphid per Petri dish, with 12 dishes in total) for 14 hours, and recorded the winged phenotype of the progeny (when they reached adulthood) produced by each group of adults (24 h post-treatment) as previously described (Parker and Brisson 2019).

Subsequently, we exposed *M. euphorbiae* aphids to high- and low-quality tomato plants (grown under light- and water-depleted conditions) as a cue to trigger the production of winged morphs (Mittler and Dadd 1966; Müller, et al. 2001; Wadley 1923). Twelve hours after treatment, we collected *M. euphorbiae* adults in pools of four and stored them in Eppendorf tubes at −80ºC until further processing. We transferred the progeny from each group of adults onto healthy tomato plants and recorded the winged phenotype of each group when they reached adulthood.

### qPCR

We macerated pools of 4 aphids with a pestle in 800 µL of TRIzol (Invitrogen; Thermo Fisher Scientific, Inc., Waltham, MA, USA). We extracted total RNA using Trizol-BCP (1-bromo-3-chloropropane; Life Technologies, Thermo Fisher Scientific, Inc., Waltham, MA, USA), with an isopropanol precipitation and ethanol wash. We eluted the RNA pellets in nuclease-free water. Then we used 1 μg of total RNA for cDNA synthesis with iScript cDNA synthesis kit (Bio-Rad Laboratories, Inc., Hercules, CA, USA). We designed quantitative PCR primers that amplified a 118 bp fragment of the expressed region of the densoviral-like genes (APNS556F 5’-GCGATTAGGAATTGGTGCGC-3’ and APNS675R 5’-AACGGTTCCAGGTCCCCTAG-3’) in *M. euphorbiae*. Reactions were run on a Bio-Rad CFX96 Real-Time System machine, with an initial step of 95ºC for 3 min and 40 cycles of 95ºC for 10 s and 60ºC for 30 s. We used three endogenous control genes (Glyceraldehyde 3-phosphate dehydrogenase (F 5’-CGGGAATTTCATTGAACGAC-3’ & R: TCCACAACACGGTTGGAGTA), NADH dehydrogenase (F 5’-CGAGGAGAACATGCTCTTAGAC-3’ & R 5’-GATAGCTTGGGCTGGACATATAG-3’), and Ribosomal Protein L32 (F 5’-CAAAGTGATCGTTATGACAAACTCAA-3’ & R 5’-CGTCTTCGGACTCTGTTGTCAA -3’)) (Chung, et al. 2020; Li, et al. 2020), which were similarly expressed in both morphs. qPCR reactions (20 μL) included 1X PCR buffer (Invitrogen, Waltham, MA, USA), Mg2+ at 2 mM, dNTPs at 0.2 mM, 1X EvaGreen (Biotium, Fremont, CA, USA), 0.025 units/mL of Invitrogen Taq, and 100 ng cDNA. Primer efficiencies were optimized to 100 ± 5% using a 10-fold serial dilution of cDNA (g3PDH: 400 nM forward & 350 nM reverse; NADH: 350 nM forward & 300 nM reverse, RpL32 400 nM for both primers, and APNS-2 homolog 200 nM for both primers). All qPCR reactions were run in triplicate. We evaluated densoviral-like gene expression in pools of four winged and four unwinged adult aphids, and in pools of four parental aphids that produced different percentages of winged and unwinged offspring 12 hours after wing induction assays. All assays were conducted in three biological replicates.

## Supplementary materials

**Table S1:** Macrosiphini genomic and transcriptomic data sets used to construct the aphid phylogeny and screen for densoviral EVEs.

**Table S2:** Densoviral genomes assembled from publicly available transcriptomes.

**Table S3:** Collection details of aphids used for sequencing.

**Table S4:** *Aphis* species genomic data sets used as the outgroup in the Macrosiphini phylogeny.

**Table S5:** Sequences and accession numbers of the major aphid densoviral proteins used for the Macrosiphini densoviral EVE screening.

**Table S6:** Sequences and accession numbers of aphid non-structural proteins used to reconstruct the Densovirus phylogeny.

**Figure S1:** Genome assembly metrics of *Neomyzus circumflexus* and *Uroleucon eupatorifoliae*.

**Figure S2:** Macrosiphini phylogeny based on a maximum-likelihood analysis of ultra-conserved elements (UCEs) from aphid species with sequenced genomes (as of January 2026). Bootstrap support values below 100 are shown on the corresponding branches and represent percentages from 500 replicates. The tree was inferred unrooted with RAxML-NG and rooted for display using Aphis as the outgroup.

## Acknowledgments

Thanks to Taylor Do for technical assistance, and to Sevier and Cocke County farmers for logistical support during aphid collections.

## Data availability

The genome assemblies generated in this study are available in NCBI under accessions ASM4999865v1 (*Uroleucon eupatorifoliae*) and ASM5896979v1 (*Neomyzus circumflexus*). The genome sequence of Neomyzus circumflexus densovirus (NcDV) is available in NCBI under accession PZ554439. Raw RNAseq reads are available in the NCBI Sequence Read Archive; accession numbers are listed in Table S1. Raw Nanopore DNA sequencing reads for *N. circumflexus* are available in the NCBI Sequence Read Archive under BioProject Accession PRJNA1442791. Bioinformatic code is available at https://github.com/Artifice120/Uroleucon-eupatorifoliae-assembly-to-annotation, https://github.com/Artifice120/Neomyzus-assembly, and https://github.com/Artifice120/UCE_Aphidae_Phylogeny-.

