## Supplementary figures and images for "Evolution of endogenized densoviral elements across aphid species"

### Figure S1

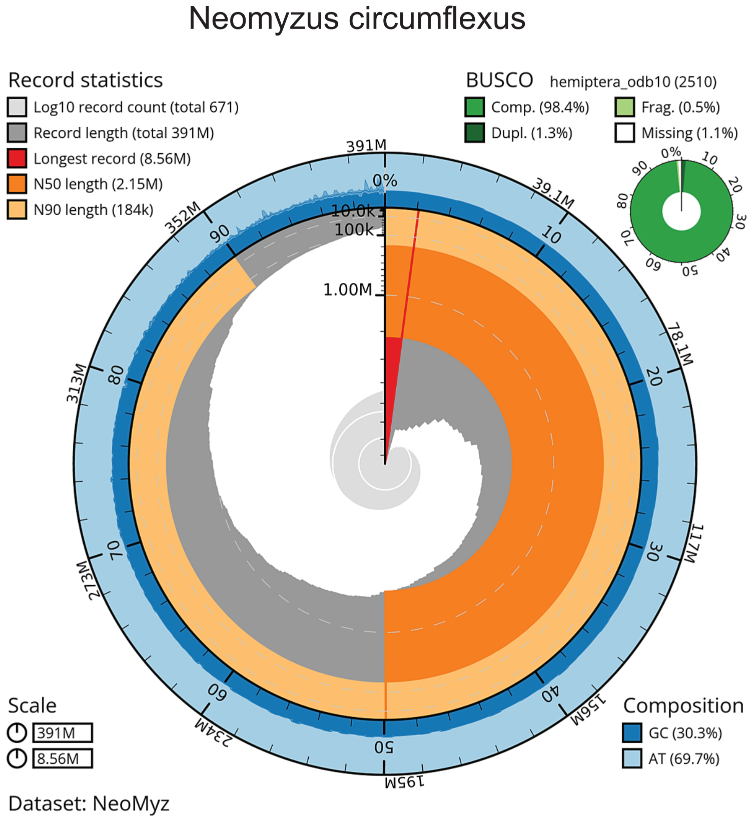

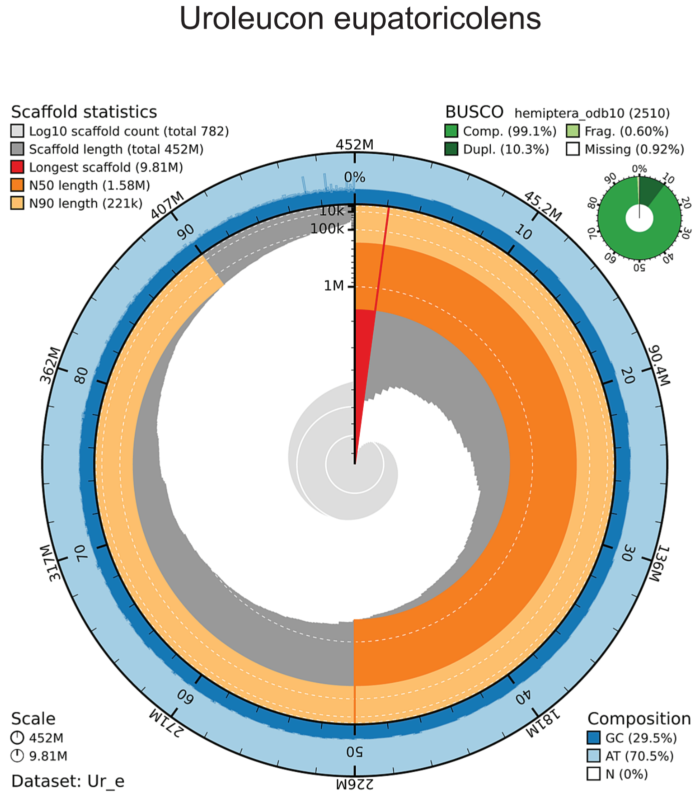

### Figure S2

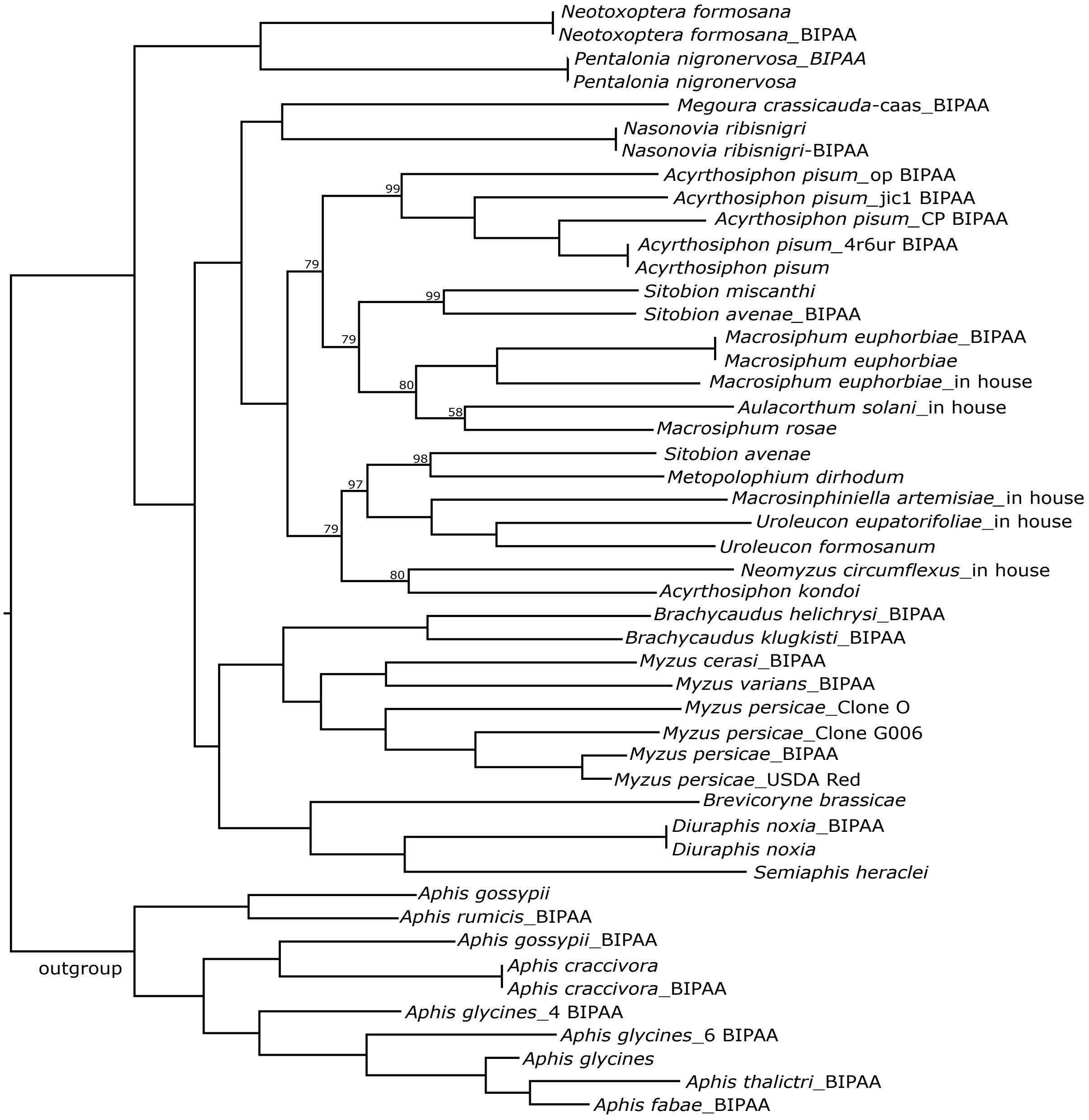
